# Characterization of immunoactive and immunotolerant CD4+ T cells in breast cancer by measuring activity of signaling pathways that determine immune cell function

**DOI:** 10.1101/2020.10.08.292557

**Authors:** Yvonne Wesseling-Rozendaal, Arie van Doorn, Karen Willard-Gallo, Anja van de Stolpe

**Author notes:** Arie van Doorn.

## Abstract

Cancer immunotolerance can be reversed by checkpoint blockade immunotherapy in some patients, but response prediction remains a challenge. CD4+ T cells play an important role in activating adaptive immune responses against cancer. Conversion to an immune suppressive state impairs the anti-cancer immune response and is mainly effected by CD4+ Treg cells. A number of signal transduction pathways activate and control functions of CD4+ T cell subsets. As previously described, assays have been developed which enable quantitative measurement of the activity of signal transduction pathways (e.g. TGFβ, NFκB, PI3K-FOXO, JAK-STAT1/2, JAK-STAT3, Notch) in a cell or tissue sample. Using these assays, pathway activity profiles for various CD4+ T cell subsets were defined and cellular mechanisms underlying breast cancer-induced immunotolerance investigated *in vitro*. Results were used to measure the immune response state in a clinical breast cancer study.

**Methods:** Signal transduction pathway activity scores were measured on Affymetrix expression microarray data of resting and immune-activated CD4+ T cells, immune-activated CD4+ T cells incubated with breast cancer tissue supernatants, CD4+ Th1, Th2, and Treg cells, and of clinical study samples in which CD4+ T cells were derived from blood, lymph node and cancer tissue from primary breast cancer patients (n=10).

**Results:** *In vitro* CD4+ T cell activation induced PI3K, NFκB, JAK-STAT1/2, and JAK-STAT3 pathway activity. Simultaneous incubation with primary cancer supernatant reduced PI3K and NFκB, and partly reduced JAK-STAT3, pathway activity, while simultaneously increasing TGFβ pathway activity; characteristic of an immune tolerant state. CD4+ Th1, Th2, and Treg cells all had a specific pathway activity profile, with activated immune suppressive Treg cells characterized by high NFκB, JAK-STAT3, TGFβ, and Notch pathway activity scores. An immune tolerant pathway profile was identified in CD4+ T cells from tumor infiltrate of a subset of primary breast cancer patients which could be contributed to activated Treg cells. A Treg pathway profile was also identified in blood samples.

**Conclusion:** Signaling pathway assays can be used to quantitatively measure the functional immune response state of lymphocyte subsets *in vitro* and *in vivo*. Clinical results suggest that in primary breast cancer the adaptive immune response of CD4+ T cells has frequently been replaced by immunosuppressive Treg cells, potentially causing resistance to checkpoint inhibition. *In vitro* study results suggest that this effect is mediated by soluble factors from cancer tissue (e.g. TGFβ). Signaling pathway activity analysis on TIL and/or blood samples is expected to improve predicting and monitoring response to checkpoint inhibitor immunotherapy.

## Introduction

Infiltration of cancer tissue by a variety of immune cells is necessary to mount an adequate immune response against the cancer cells. During the past decades, evidence has been accumulating that cancer tissue can be highly successful in creating an immune-tolerant environment, which interferes with the appropriate anti-cancer immune response of tumor infiltrating T cells. Checkpoint inhibitor immunotherapy against cytotoxic T lymphocyte antigen-4 (CTLA-4) and Programmed Death-1 (PD-1) aims at restoring the effector function of CD8+ cytotoxic T cells, and when successful, has been shown to have the potential of being a curative treatment (1). An increasing number of immunotherapy drugs that aim at re-activating the immune response in such a targeted manner is in development. Unfortunately, only a limited percentage of patients respond to this type of immunotherapy (2). Assays to predict response, such as PD-1/PDL-1 and CD4+/CD8+ immunohistochemistry (IHC) staining measurements have proven to be not sufficiently reliable in predicting response in the individual patient. Consequently, there is a high need for assays to predict therapy response and to assess, as soon as possible, whether the installed therapy is effective (2–5).

In the tumor infiltrate, CD4+ T cells play an important role in activating adaptive immune responses, and when reverted to an immune suppressed state they impair the anti-cancer immune response. Their functional state is regulated by signal transduction pathways, e.g. the PI3K, JAK-STAT1/2, JAK-STAT3, NFκB, TGFβ, and Notch pathways (6–11). Recently, assays have been developed which enable quantitative measurement of the activity of these signal transduction pathways in tissue and blood samples (12–16).

These signaling pathway assays were used to *in vitro* characterize resting, immune-activated and immunotolerant CD4+ T cells, as well as CD4+ T-helper 1 (Th1), T-helper 2 (Th2), and regulatory T (Treg) cells, in terms of the activity of signaling pathways. We show that the identified pathway activity profiles can be of help in elucidating the mechanism leading to cancer-induced immunotolerance, and to investigate the type and the functional state of CD4+ T cells in blood, lymph node, and tumor infiltrate (TIL) samples from cancer patients (17).

## Methods

### Signaling pathway activity analysis of preclinical and clinical studies

Tests to quantitatively measure functional activity of PI3K, NFκB, TGFβ, JAK-STAT, and Notch signal transduction pathways on Affymetrix Human Genome U133 Plus 2.0 expression microarrays data have been described before (13, 14, 16, 18, 19). In brief, the pathway assays are based on the concept of a Bayesian network computational model which calculates from mRNA levels of a selected set, usually between 20 and 30, target genes of the pathway-associated transcription factor a probability score for pathway activity, which is translated to a log2 value of the transcription factor odds, i.e. a log2odds score scaling with pathway activity. The models have all been calibrated on a single cell type and were subsequently frozen and validated on a number of other cell and tissue types without further adaptations of the models. The range (minimum-maximum pathway activity) on the log2odds scale is different for each signaling pathway. While the signaling pathway assays can be used on all cell types, the log2odds score range may vary per cell/tissue type. Of note, measurement of the activity of the PI3K pathway assay is based on the inverse inference of activity of the PI3K pathway from the measured activity of the FOXO transcription factor, in the absence of cellular oxidative stress (13). For this reason, the FOXO activity score is presented in the figures instead of PI3K pathway activity. In cell culture experiments *in vitro*, PI3K pathway activity can be directly (inversely) inferred from FOXO activity. Activity scores were calculated for the FOXO transcription factor, and NFκB, JAK-STAT, Notch and TGFβ signaling pathways on Affymetrix expression microarray datasets from the Gene Expression Omnibus (GEO, www.ncbi.nlm.nih.gov/geo) database (20) or generated in our own lab, and presented on a log2odds scale (12–14).

The GSE71566 dataset contained Affymetrix data from CD4+ T cells isolated from cord blood. Naive CD4+ T cells were compared with CD4+ T cells which had been activated by treatment with anti-CD3 and anti-CD28, and with CD4+ T cells which had been differentiated to either Th1 or Th2 cells (21). The GSE11292 dataset contained data from cell-sorted Treg cells (CD4+CD25hi), activated *in vitro* with anti-CD3/CD28 in combination with interleukin 2 (IL-2) in a time series (22).

Two Affymetrix datasets had been generated in our own lab and have been described in detail before (17) and are also publically available under accession numbers GSE36765 and GSE36766. The GSE36766 dataset contains Affymetrix data from an *in vitro* study in which breast cancer tissue sections from fresh surgical specimens of primary untreated breast cancers (n=4) were mechanically dissociated in X-VIVO 20 medium; CD4+ T cells from healthy donor blood were incubated for 24 hours with anti-CD3/CD28 with or without primary tumor X-VIVO 20 supernatant, and microarray analysis was performed. The GSE36765 dataset is a clinical dataset containing data from patients with primary untreated breast cancer; methods and patient characteristics have been described by us in detail before (17). In brief, CD4+ T cells were isolated from primary tumors, axillary lymph nodes, and peripheral blood of ten patients with invasive breast carcinomas, and peripheral blood of four healthy donors.

### Microarray data source and quality control

Analyzed datasets contained Affymetrix Human Genome U133 Plus 2.0 expression microarray data. Quality control (QC) was performed on Affymetrix data of each individual sample based on twelve different quality parameters following Affymetrix recommendations and previously published literature (23, 24). In summary, these parameters include the average value of all probe intensities, presence of negative or extremely high (> 16-bit) intensity values, poly-A RNA (sample preparation spike-ins) and labelled cRNA (hybridization spike ins) controls, *GAPDH* and *ACTB* 3’/5’ ratio, the centre of intensity and values of positive and negative border controls determined by affyQCReport package in R, and an RNA degradation value determined by the AffyRNAdeg function from the Affymetrix package in R (25, 26). Samples that failed QC were removed prior to data analysis.

### Statistics

Mann-Whitney U testing was used to compare pathway activity scores across groups. In case another statistical method was more appropriate due to the content of a specific dataset, this is indicated in the legend of the figure. For pathway correlation statistics, Pearson correlation tests were performed. Exact p-values are indicated in the figures.

### Defining a threshold for abnormal pathway activity in blood CD4+ T cell samples

To enable clinical use of the pathway activity test on blood samples a preliminary threshold value was calculated above which signaling pathway activity may be considered as abnormally high. The GSE36765 dataset contained blood derived CD4+ T cell samples from four healthy individuals. For each pathway, the mean pathway activity score ± 2SD was calculated and the upper threshold for normal defined as the mean+2SD.

## Results

The activity of the FOXO transcription factor, and of the NFκB, JAK-STAT1/2, JAK-STAT3, Notch, and TGFβ signaling pathways was measured in resting and activated CD4+ T cells, in CD4+ T cells differentiated to Th1 and Th2 cells, in Treg cells, in CD4+ T cells incubated with breast cancer supernatant, and in a series of matched blood, lymph node and tumor infiltrate samples from patients with breast cancer. As mentioned, pathway activity scores are presented on a log2odds scale, with a varying activity range (from minimum to maximum measured pathway activity) on this scale per pathway and cell/tissue type.

### Signaling pathway activity in resting and activated CD4+ T cells, in CD4+ derived Th1 and Th2, and in CD4+ Treg cells

CD4+ T cells can have different phenotypes with specialized functions in the immune response, such as Th1, Th2, and the strongly immune-suppressive Treg cells. We performed signaling pathway analysis on these different CD4+ T cell types generated *in vitro*, and observed very specific signaling pathway activity profiles for the different cell types. In all samples, PI3K pathway activity could be inversely inferred from FOXO activity since there was no evidence for cellular oxidative stress interfering with the inverse relationship between FOXO and PI3K pathway activity (results not shown) (13). Conventional activation of CD4+ T cells obtained from cord blood with anti-CD3/anti-CD28 antibodies resulted in activation of the NFκB pathway, and slightly increased JAK-STAT1/2 and JAK-STAT3 pathway activity, while the activity of the TGFβ pathway, already low, was further reduced (Figure 1A). FOXO activity decreased, indicating activation of the PI3K growth factor pathway. Activated T cells which had been further differentiated *in vitro* to respectively Th1 and Th2 T cells, showed differential changes in pathway activity: NFκB and JAK-STAT1/2 pathway activity increased in Th1, while the activity of the FOXO transcription factor increased in Th2 cells (Figure 1B).

**Figure 1.**
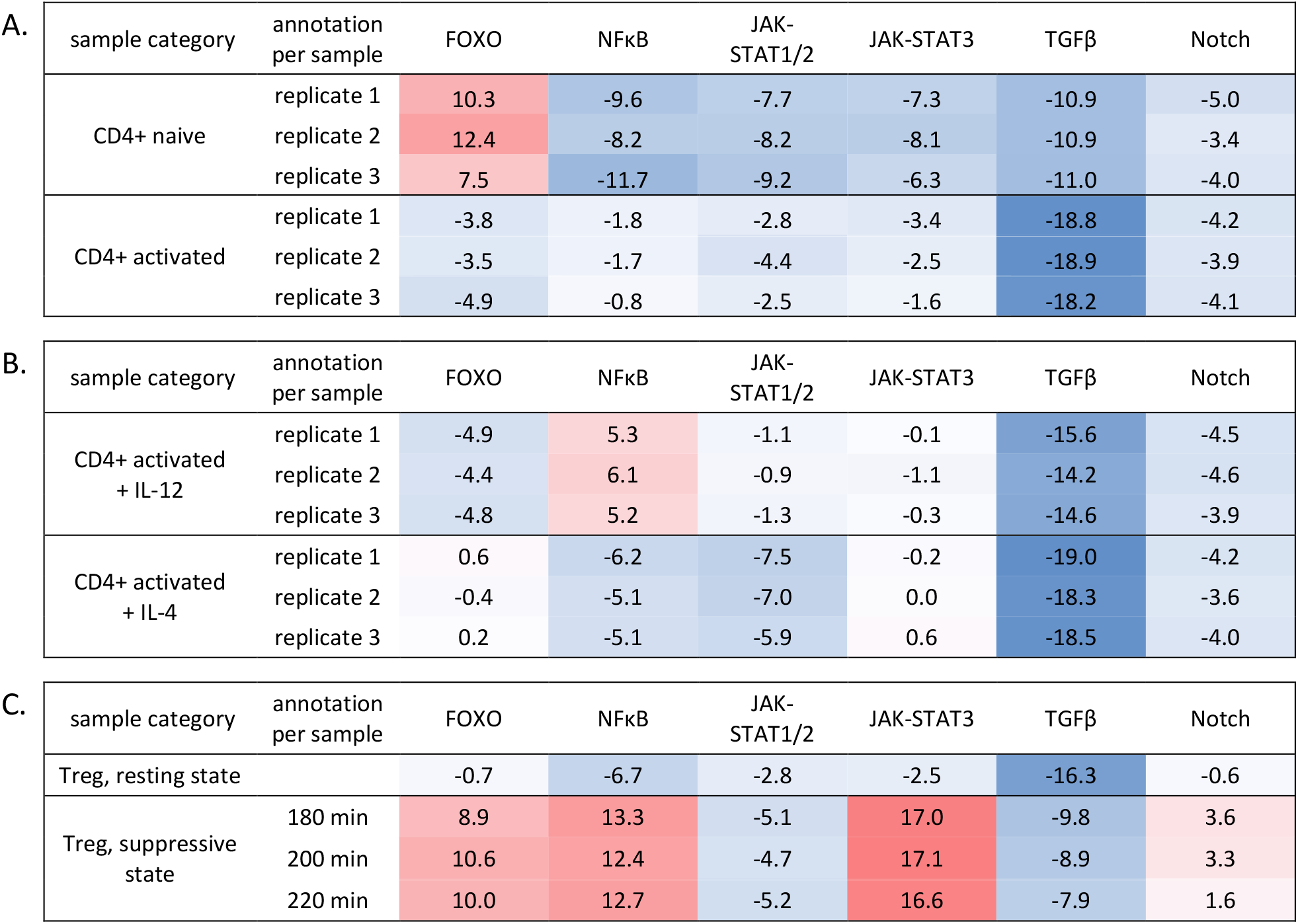
Signal transduction pathway activity in CD4+ T cells. A,B. GSE71566, CD4+ T cells derived from cord blood, resting or activated using anti CD3/CD28 (A), or stimulated with IL-12 and IL-4 for differentiation towards respectively Th1 and Th2 T cells (B) (21). C. GSE11292, CD4+ Treg cells, resting or activated with anti-CD3/-CD28 with IL-2 (22). The first column contains the sample annotation as available from GEO; top of columns contains names of signaling pathways; note that FOXO transcription factor activity is presented which is the inverse of PI3K pathway activity. Signaling pathway activity scores are presented on log2odds scale with color coding ranging from blue (most inactive) to red (most active).

In activated CD4+ T cells from our own dataset, in which CD4+ T cells had been derived from peripheral blood from healthy volunteers and subsequently had been activated by anti-CD3/anti-CD28 incubation, a similar CD4+ T cell pathway activity profile was observed after activation with anti-CD3/CD28 (Figure 2, compare not activated-control with activated-control) (17).

**Figure 2.**
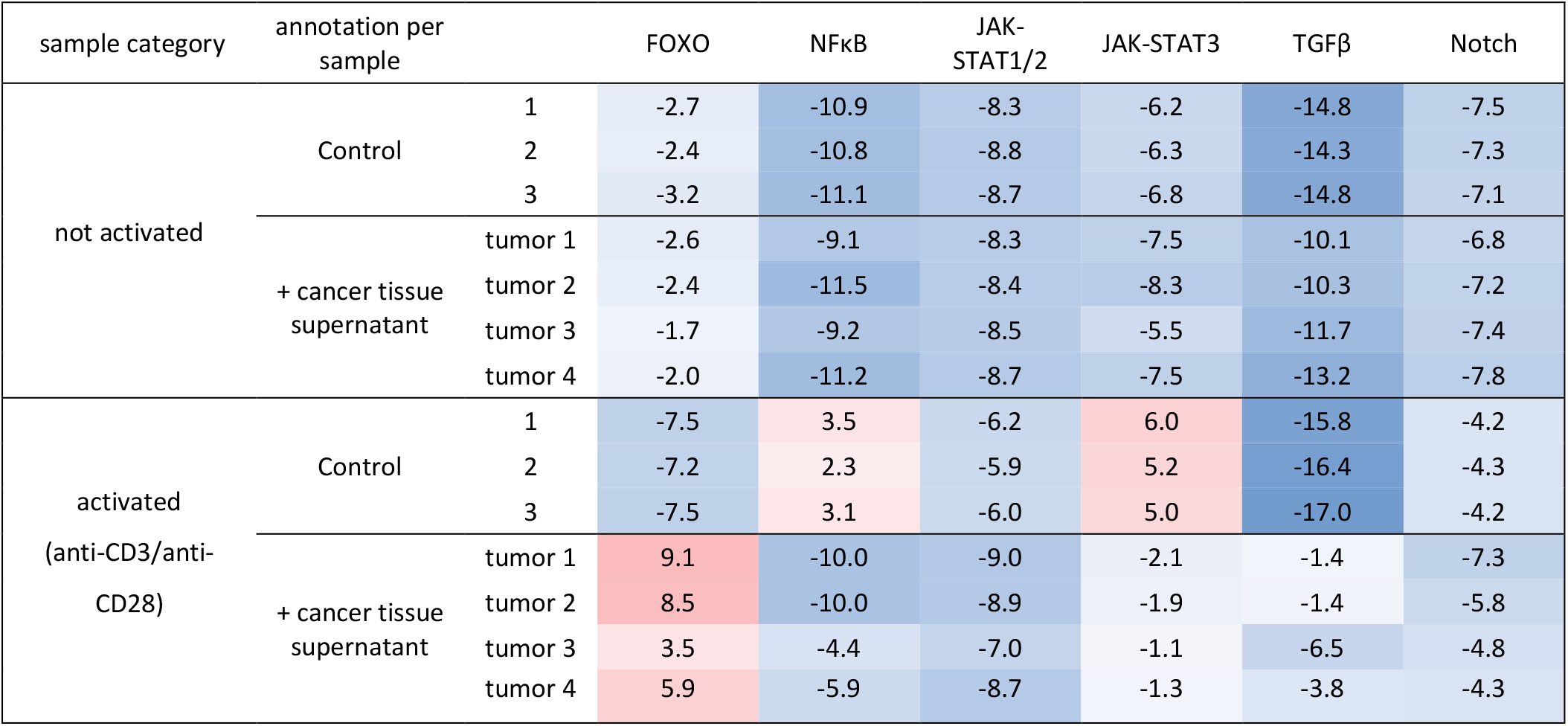
Effect of breast cancer tissue supernatant on signal transduction pathway activity scores in resting and activated CD4+ T cells. Dataset GSE36766, resting or in vitro activated (anti-CD3/CD28) CD4+ T cells from healthy donor blood were incubated with primary cancer tissue supernatant (n=4 different cancers) or control medium (n=3) (17). The left set of columns contain information on the experimental protocol for each sample. Signaling pathway activity scores are presented on a log2odds scale with color coding ranging from blue (most inactive) to red (most active).

A distinct type of CD4+ T cells are the Treg cells, which in the analyzed set had been activated with anti-CD3/CD28 plus IL-2. Treg cells were clearly distinguishable from the other CD4+ T cell types. Resting state Treg cells mostly resembled Th2 cells, but with a slightly higher JAK-STAT1/2, lower JAK-STAT3, and higher Notch pathway activity (Figure 1C). Induction of the Treg immune suppressive state (induced Treg cells; iTreg) was associated with increased FOXO activity, reflecting a reduction in PI3K pathway activity, increased NFκB, JAK-STAT3, TGFβ, and Notch pathway activity, and reduced JAK-STAT1/2 pathway activity (Figure 1C). These quantitative pathway analysis results support a mechanistic role for the PI3K-FOXO, NFκB, JAK-STAT1/2 and JAK-STAT3, pathways during activation of CD4+ T cells differentiation to Th1 and Th2 T cells, while the activity of the TGFβ and Notch signaling pathways was specifically linked to the iTreg cells.

### Immune suppressive effect of cancer cell supernatant on activated CD4+ T cells

Our previously published study in which we investigated the effect of adding breast cancer tissue supernatant (SN) to resting or activated CD4+ T cells of healthy individuals, allowed us to analyze the effect of breast cancer tissue on signaling pathway activity in CD4+ T cells (Figure 2). While the addition of supernatants of four different dissected primary breast cancer tissue samples did not affect signaling pathway activities in resting CD4+ T cells, the same supernatants changed pathway activity scores in activated CD4+ T cells: NFκB, and JAK-STAT1/2 and JAK-STAT3 pathway activity decreased, while the activity of the TGFβ pathway increased (Figure 2). In addition, FOXO activity increased, reflecting a decrease in PI3K pathway activity as a consequence of incubation with tumor supernatant. Importantly, after the addition of SN, the JAK-STAT3 pathway activity remained higher than in resting CD4+ T cells, while the activity scores of the FOXO transcription factor increased even above those seen in resting CD4+ T cells. In summary, cancer tissue supernatant caused a partial reversion of the pathway activity profile towards that of resting CD4+ T cells in combination with a strong upregulation of the immune-suppressive TGFβ pathway and decrease in PI3K pathway activity, in line with the induction of an immunotolerant state.

### Clinical stud: signaling pathway activity in CD4+ T cells derived from blood, lymph node, and breast cancer tissue samples of patients with primary breast cancer

Having characterized the signaling pathway activity profile associated with activated (anti-CD3/CD28) and immunotolerant CD4+ T cells *in vitro*, we proceeded to investigate CD4+ T cells that had been obtained previously in a clinical patient setting (17). Matched CD4+ T cells from peripheral blood, lymph nodes, and tumor infiltrate of ten early untreated breast cancer patients were compared with respect to the above identified signaling pathway activities.

Notch pathway activity was significantly increased (p=0.002) in blood samples from breast cancer patients compared to healthy individuals (Figure 3). Increased JAK-STAT3, and TGFβ pathway activity scores were measured in a sample subset, but compared to the four control samples this did not reach a statistically significant difference (Figure 3). In CD4+ T cells isolated from tumor infiltrate (TIL), NFκB, JAK-STAT1/2, JAK-STAT3, TGFβ, and Notch pathway activity were significantly increased compared to the pathway activity scores in corresponding patient blood samples. FOXO activity was also increased in TIL, indicative of an inactivated PI3K pathway (Figure 3; Supplementary Figure S1). Comparing absolute pathway activity scores of TIL samples with the pathway scores measured in the *in vitro* cancer supernatant study, TIL-derived CD4+ T cells mostly resembled the *in vitro* activated CD4+ T cells that had been incubated with cancer tissue supernatant (Figures 2,3). To identify which subset of CD4+ T cells was responsible for the observed pathway profile in the TIL, this was compared with the *in vitro* identified profiles. The TIL-derived CD4+ T cells profile was in the majority of patients highly similar to the iTreg cell profile, characterized by low PI3K pathway activity, high JAK-STAT3, NFκB, TGFβ and Notch pathway activity.

**Figure 3.**
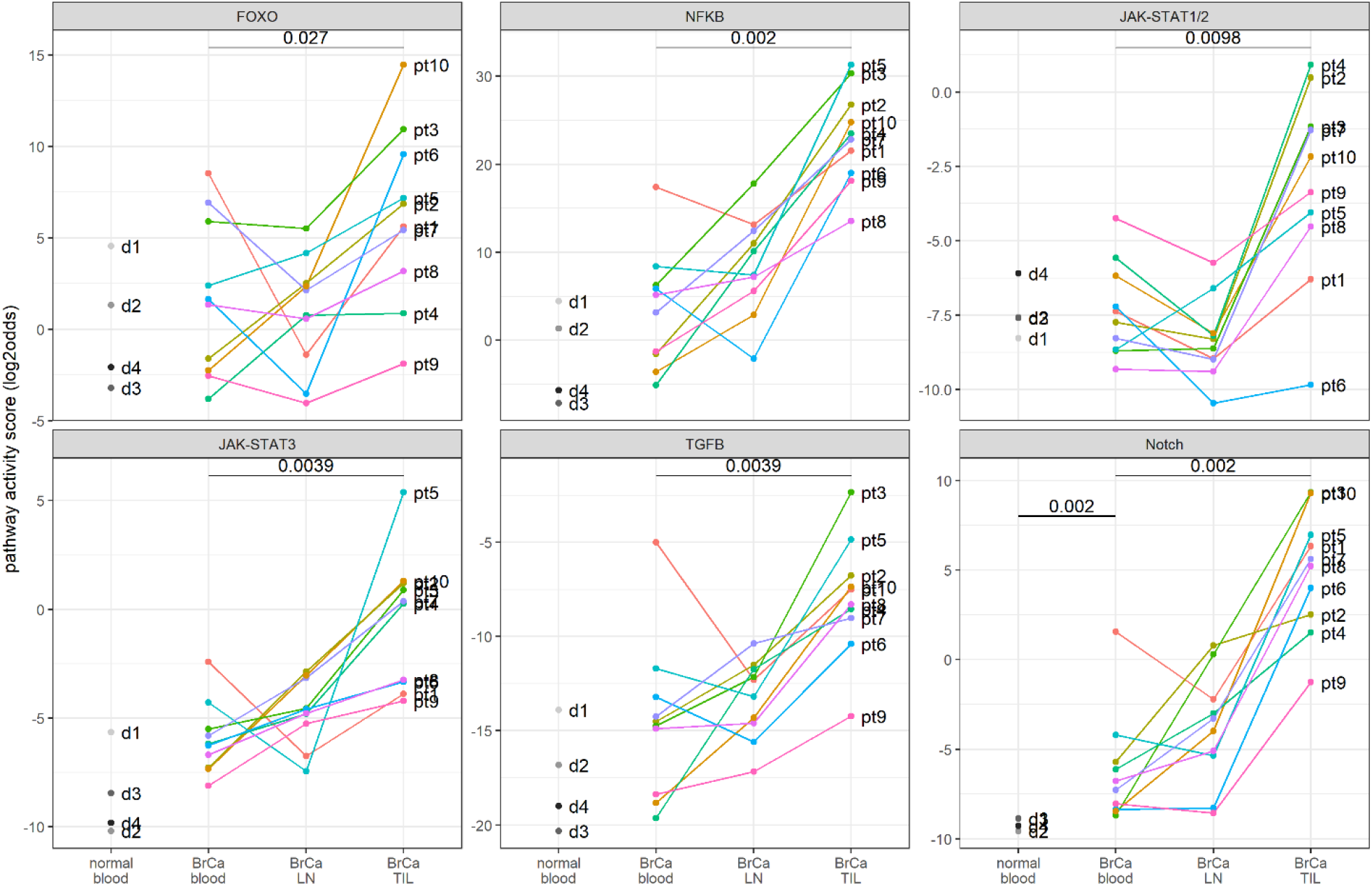
Signal transduction pathway activity in CD4+ T cells from matched blood, lymph node (LN), and TIL samples of ten patients with invasive breast cancer (BrCa) and four healthy donors; dataset GSE36765 (17). From left to right: FOXO, NFκB, JAK-STAT1/2, JAK-STAT3, TGFβ, and Notch pathway activity scores. Pathway activity is indicated as log2odds on the y-axis. Individual sample pathway scores are matched by color-coding (‘pt’ for BrCa patient samples; ‘d’ for healthy donor samples). P-values (two-sided Wilcoxon-test) are listed when significant (p<0.05); a paired test was performed when comparing amongst BrCa samples.

The activity of the NFκB, JAK-STAT3, TGFβ, and Notch signaling pathways was strongly correlated in TIL samples, suggesting that they were often combined active in the same CD4+ T cells, and that these CD4+ T cells are iTreg cells (Supplementary Figure S2).

Compared to the *in vitro* T cell experiments, in the clinical samples pathway activity scores varied more between patients, reflecting heterogeneity of the analyzed breast cancer patient group that consisted of four ER/PR positive luminal patients, one ER/PR/HER2 positive patient, and five triple negative patients (17) (Figure 4; Supplementary Figure S1). Although the number of patients per subtype was too small to draw any conclusions on subtype specific pathway profiles in CD4+ TIL cells, it is interesting to note that TGFβ and NFκB pathway activity, were significantly lower in triple negative patients compared to other subtypes (p=0.028 and p=0.028 respectively). In contrast, in ER/PR positive (HER2 negative) cancer samples the immune suppressive iTreg profile (high NFκB, JAK-STAT3, TGFβ, and Notch pathway activity scores) was most prominently present, compared to the triple negative cancers.

**Figure 4.**
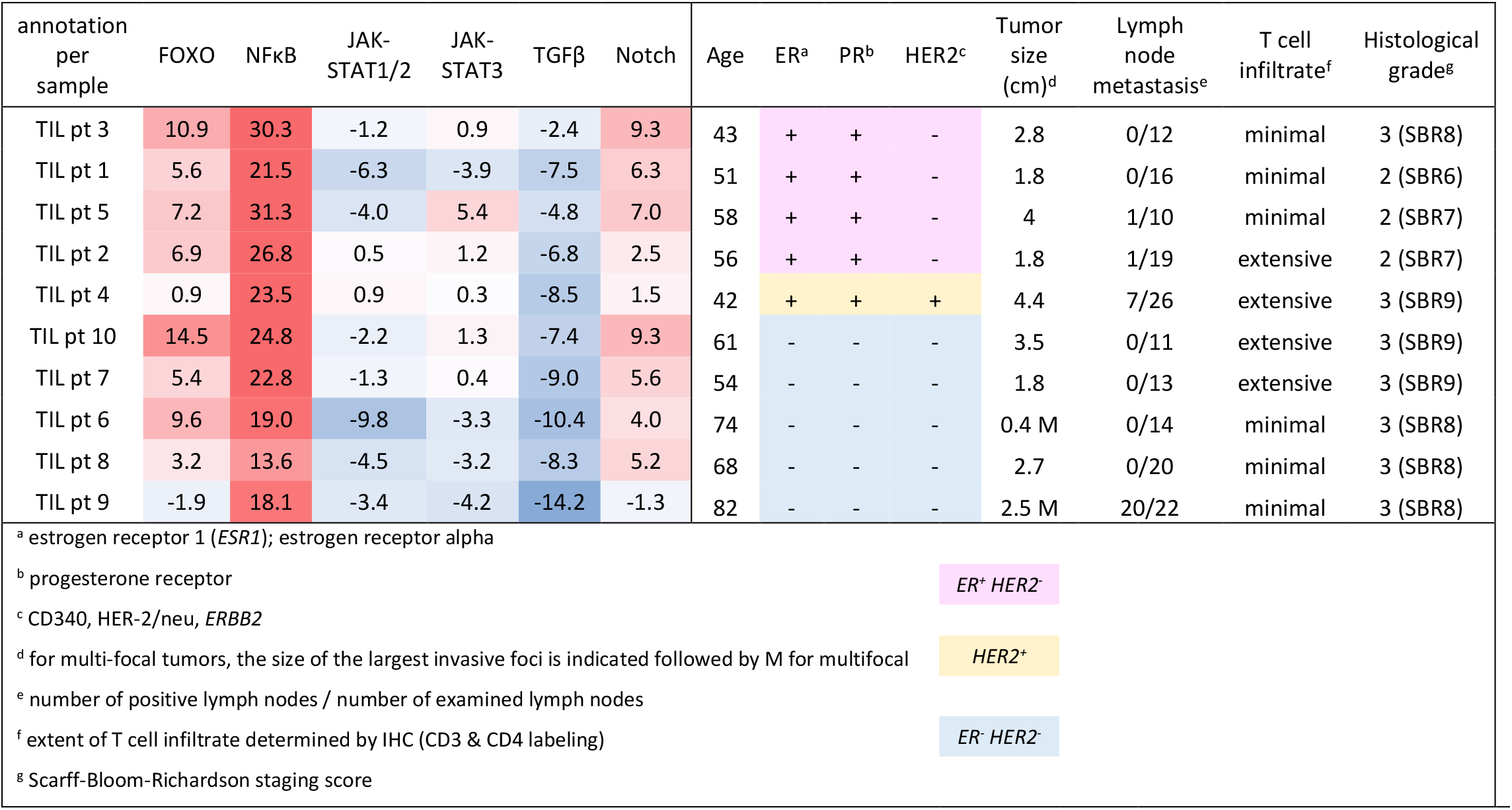
GSE36765. Pathway activity in TIL CD4+ T cells, related to the additional clinical information which was described before (17). Signaling pathway activity scores are presented as log2odds with color coding ranging from blue (most inactive) to red (most active).

### A threshold for abnormal pathway activity in blood CD4+ T cell samples

For each pathway, the mean pathway activity score ± 2 standard deviations (SD) was calculated and the upper threshold for normal defined as the mean + 2SD (Supplementary Table 1). When applying these thresholds to the patient samples, combined Notch, JAK-STAT3, and TGFβ pathway activity was identified in 2 patients, abnormal Notch pathway activity in 9 out of 10 patients (Table 1, and Supplementary Table S1).

**Table 1.**
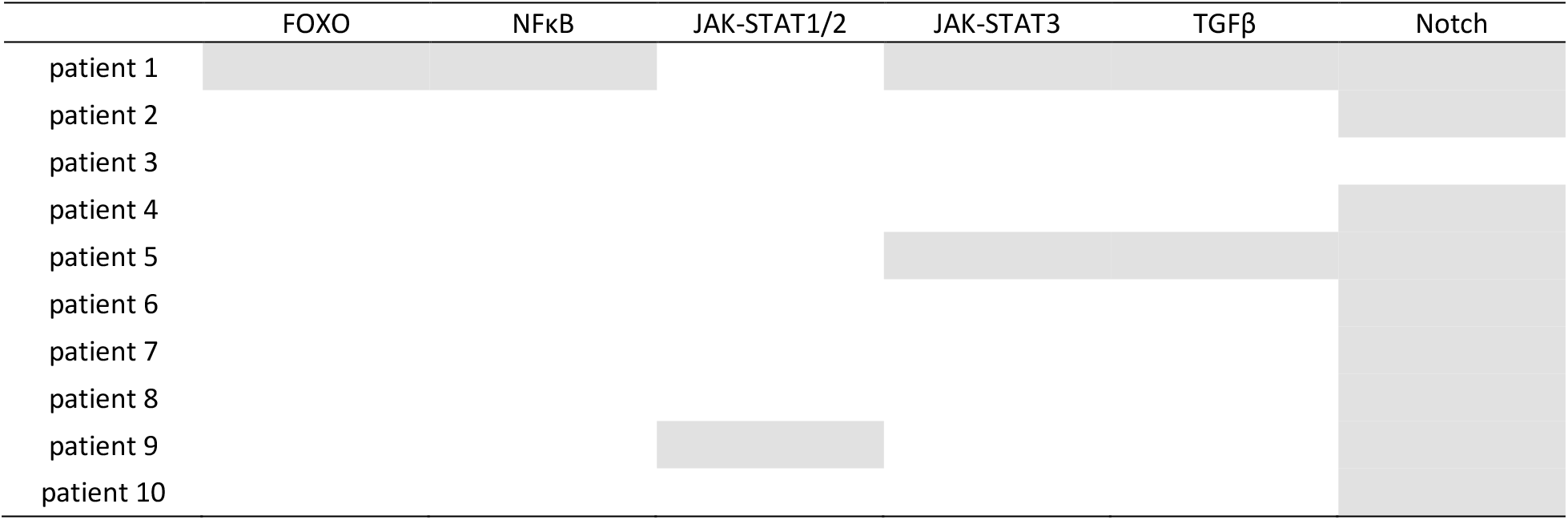
Abnormal pathway activities in blood derived CD4+ T cells from ten patients with primary breast cancer (GSE36765, (17)), using normal pathway activity score + 2SD as an upper threshold for normal.

## Discussion

The recently developed assay platform for measuring the activity of the most relevant signal transduction pathways was used to *in vitro* characterize the functional state of different subsets of CD4+ T cells with respect to signaling pathway activity, identify the pathway profile of immunotolerant CD4+ T cells, and analyze the immunosuppressive effect of cancer tissue supernatant. Subsequently, the comparison was made with CD4+ T cell samples from breast cancer patients to identify the CD4+ T cell subset responsible for immunotolerance in TIL, and to investigate whether the immunotolerant CD4+ T cell pathway profile can also be detected in blood samples from breast cancer patients. The signal transduction pathway assays used in the current study have been biologically validated on multiple cell types, including immune cell types (12– 14, 16). Affymetrix expression microarray data were used for the signaling pathway analysis, however, it is good to keep in mind that with this assay technology Affymetrix microarrays are only used as a method to measure a carefully preselected set of mRNA levels, and not to discover a new gene profile. From Affymetrix whole transcriptome data, expression data of only 20-30 pathway target genes per signaling pathway are used to calculate a signaling pathway activity score, the respective genes have been described (12–14, 16, 18).

The measured changes in pathway activity scores associated with CD4+ T cell activation were consistent across two independent studies (17, 21). Increased PI3K (inferred from a decrease in FOXO activity), NFκB, JAK-STAT1/2, and JAK-STAT3 pathway activity reflects T cell activation, and is of crucial importance for clonal proliferation and execution of specific immune response functions (6–8, 27–32). Differentiation of activated CD4+ T cells to Th1, Th2 and Treg cells appeared to be associated with characteristic alterations in the pathway activity profile, reflecting specific cell functions determined by signaling pathway activity (13, 14, 16, 19). For example, In Th1 compared to Th2 cells, a higher JAK-STAT1/2 pathway activity is necessary for the Th1 role to activate CD8+ T cells, for example in viral infections (7), while a high PI3K pathway activity is required for expression of the TBET transcription factor, essential for Th1 function (6). In immune suppressive iTreg cells, the identified combined activity of Notch with JAK-STAT3 and TGFβ pathways is in line with the Notch pathway controlling the immune suppressive function in cooperation with the other signaling pathways (33, 34). The very low PI3K pathway activity reflects the minimal role for this signaling pathway in Treg cells (6).

Cancer tissue supernatant appeared to induce a partial reversal of the CD4+ activation profile *in vitro*, resulting in a profile which was very similar to that described for immunotolerant lymphocytes and suggesting that one or more soluble factors from cancer tissue had induced immunotolerance (31, 35, 36). An earlier analysis of this study using conventional bioinformatics approaches had resulted in a similar conclusion as to a tumor suppressive effect of cancer supernatant, but without linking this to signaling pathway activity (17). Potential candidates capable of inducing the identified pathway activity profile are a soluble form of PD-L1, TGFβ, and IL-6 (9, 37–43). The PD-L1 receptor PD-1 mediates tolerance induction in part by inhibiting activation of PI3K, explaining the observed inhibition of the PI3K pathway (36, 44). TGFβ is locally produced and can directly activate the tumor suppressive TGFβ pathway (9, 41, 42). Interleukin 6 (IL-6) induces activation of the JAK-STAT3 pathway (43). Crosstalk between these signaling pathways may be required to induce full T cell tolerance (45–47).

While the JAK-STAT3 pathway was found moderately activated in immunotolerant CD4+ T cells, higher pathway activation scores were found in properly activated CD4+ T cells suggesting a dual function for this pathway. Indeed, the transcriptional program of the STAT3 transcription factor depends on crosstalk with other signaling pathways in the cell, resulting in T cell activation in the presence of CD3/CD28 stimulation or in immunotolerance in the presence of immunosuppressive factors like TGFβ, IL-6, and PD-L1 (7, 9, 27, 35, 41).

Following *in vitro* CD4+ T cell subset analysis, CD4+ T cells from blood, lymphocyte, and TIL samples from patients with untreated primary breast cancer were analyzed. Variation in pathway activity scores between patients was larger in these clinical samples, probably caused by variable immune responses determined by factors such as antigenicity of the tumor and genetic variations.

A subset of TIL-derived samples, mostly from luminal (ER/PR positive) breast cancers, had a pathway activity profile resembling the immune tolerant pathway profile found in the tumor supernatant-treated cells with respect to PI3K, JAK-STAT3, and TGFβ pathway activity, complemented by higher Notch and NFκB pathway activity. Comparison of these TIL samples with Th and Treg subset pathway profiles revealed a strong match with iTreg cells, including Treg-specific Notch pathway activity. Treg cells induce immune suppression in cancer tissue and are thought to mediate resistance against checkpoint inhibitors (48–50). Indeed patients with ER/PR positive breast cancer are generally unresponsive to checkpoint inhibitor treatment (51). The higher JAK-STAT1/2 and NFκB pathway activities in TIL compared to *in vitro* iTreg cells may be due to local inflammatory conditions in cancer tissue (52).

In patient blood CD4+ T cells, Notch pathway activity was significantly increased compared to healthy controls, similarly indicative of the presence of immune suppressive Treg cells. JAK-STAT3 and TGFβ pathway activity were also higher in patient blood samples but this did not reach significance, which can be explained by a mixture of CD4+ T cell subsets being present in blood and a smaller fraction of activated Treg cells than in the TIL (48). Using bioinformatics analysis, in the earlier reported analysis no significant differences in the transcriptome had been identified between CD4+ T cells from healthy and cancer patients (17). Current findings may be explained by a Treg subset of CD4+ T cells having cycled from cancer tissue into blood; alternatively, elevated levels of circulating cytokines like IL-6, IL-10 and TGFβ may explain the tumor suppressive profile in these blood samples (48, 53).

It is of potential clinical relevance that already in primary untreated breast cancer patients activated immune suppressive Treg cells pathway can be present in peripheral blood. While Treg cells can be identified and isolated using flow cytometry and FACS techniques, our pathway analysis enables assessment of resting versus activated functional state. The first clinical study is in progress to investigate whether measuring this activated Treg profile in blood may help to predict resistance to checkpoint inhibitor therapy. For Affymetrix-based signaling pathway analysis, the here defined (preliminary) thresholds for abnormal pathway activity will be used; thresholds will be adapted for qPCR-based pathway activity profiling.

### Summary and conclusion

Using Philips signaling pathway analysis, we provide evidence that soluble factors in primary breast cancer tissue induce an immune tolerant CD4+ T cell state, likely caused by activated immune suppressive Treg cells that may already be detectable in blood samples. Since activated Treg cells may confer resistance to checkpoint inhibitors, signaling pathway analysis of CD4+ T cells in TIL or in blood samples of cancer patients may enable improved prediction and monitoring of response to checkpoint inhibitor therapy.

