## Supplementary Information for "Characterization of immunoactive and immunotolerant CD4+ T cells in breast cancer by measuring activity of signaling pathways that determine immune cell function"

| sample category | annotation per sample | FOXO | NFκB | JAK-STAT1/2 | JAK-STAT3 | TGFβ | Notch | ER | WNT |
|---|---|---|---|---|---|---|---|---|---|
| CD4+ T cells from peripheral blood | donor 1 | 4.5 | 4.5 | -8.3 | -5.7 | -13.9 | -8.9 | -18.7 | -19.1 |
| | donor 2 | 1.3 | 1.4 | -7.6 | -10.2 | -16.8 | -9.6 | -19.1 | -21.0 |
| | donor 3 | -3.2 | -7.1 | -7.6 | -8.4 | -20.3 | -8.9 | -18.9 | -24.5 |
| | donor 4 | -2.1 | -5.7 | -6.1 | -9.8 | -19.0 | -9.3 | -18.9 | -21.4 |
| | patient 1 | 8.5 | 17.4 | -7.4 | -2.4 | -5.0 | 1.6 | -16.0 | -20.0 |
| | patient 2 | -1.6 | -1.5 | -7.7 | -7.3 | -14.5 | -5.7 | -18.6 | -16.7 |
| | patient 3 | 5.9 | 6.3 | -8.7 | -5.5 | -14.7 | -8.7 | -18.8 | -22.8 |
| | patient 4 | -3.8 | -5.1 | -5.6 | -6.2 | -19.6 | -6.1 | -18.6 | -21.1 |
| | patient 5 | 2.4 | 8.4 | -8.7 | -4.3 | -11.7 | -4.2 | -18.2 | -15.9 |
| | patient 6 | 1.7 | 5.9 | -7.2 | -6.3 | -13.2 | -8.4 | -18.8 | -21.2 |
| | patient 7 | 6.9 | 3.2 | -8.3 | -5.8 | -14.3 | -7.3 | -18.9 | -19.2 |
| | patient 8 | 1.3 | 5.2 | -9.3 | -6.7 | -14.9 | -6.8 | -18.7 | -17.0 |
| | patient 9 | -2.5 | -1.3 | -4.2 | -8.1 | -18.4 | -8.0 | -19.9 | -22.2 |
| | patient 10 | -2.2 | -3.6 | -6.2 | -7.3 | -18.8 | -8.4 | -19.3 | -21.8 |
| CD4+ T cells from axillary lymph nodes | patient 1 | -1.4 | 13.1 | -9.0 | -6.7 | -12.3 | -2.2 | -18.8 | -22.7 |
| | patient 2 | 2.5 | 11.0 | -8.3 | -2.9 | -11.5 | 0.8 | -17.5 | -20.8 |
| | patient 3 | 5.5 | 17.8 | -8.6 | -4.5 | -12.2 | 0.3 | -18.2 | -20.5 |
| | patient 4 | 0.8 | 10.1 | -8.2 | -4.8 | -11.7 | -3.0 | -18.2 | -21.7 |
| | patient 5 | 4.2 | 7.4 | -6.6 | -7.4 | -13.2 | -5.4 | -18.7 | -20.7 |
| | patient 6 | -3.5 | -2.1 | -10.5 | -4.6 | -15.6 | -8.3 | -19.9 | -23.4 |
| | patient 7 | 2.1 | 12.4 | -9.0 | -3.1 | -10.4 | -3.3 | -18.2 | -20.3 |
| | patient 8 | 0.6 | 7.2 | -9.4 | -4.8 | -14.6 | -5.1 | -18.7 | -26.9 |
| | patient 9 | -4.0 | 5.6 | -5.7 | -5.2 | -17.2 | -8.6 | -19.1 | -25.0 |
| | patient 10 | 2.3 | 2.9 | -8.1 | -3.0 | -14.3 | -4.0 | -19.1 | -21.9 |
| CD4+ T cells from breast tumor | patient 1 | 5.6 | 21.5 | -6.3 | -3.9 | -7.5 | 6.3 | -18.5 | -21.5 |
| | patient 2 | 6.9 | 26.8 | 0.5 | 1.2 | -6.8 | 2.5 | -15.9 | -17.5 |
| | patient 3 | 10.9 | 30.3 | -1.2 | 0.9 | -2.4 | 9.3 | -16.9 | -15.3 |
| | patient 4 | 0.9 | 23.5 | 0.9 | 0.3 | -8.5 | 1.5 | -15.7 | -19.9 |
| | patient 5 | 7.2 | 31.3 | -4.0 | 5.4 | -4.8 | 7.0 | -6.7 | -14.2 |
| | patient 6 | 9.6 | 19.0 | -9.8 | -3.3 | -10.4 | 4.0 | -19.3 | -16.6 |
| | patient 7 | 5.4 | 22.8 | -1.3 | 0.4 | -9.0 | 5.6 | -18.3 | -13.5 |
| | patient 8 | 3.2 | 13.6 | -4.5 | -3.2 | -8.3 | 5.2 | -17.5 | -25.4 |
| | patient 9 | -1.9 | 18.1 | -3.4 | -4.2 | -14.2 | -1.3 | -17.6 | -25.0 |
| | patient 10 | 14.5 | 24.8 | -2.2 | 1.3 | -7.4 | 9.3 | -18.2 | -19.1 |

*Supplementary Figure S1. Signaling pathway activities per patient from dataset GSE36765 [17].*

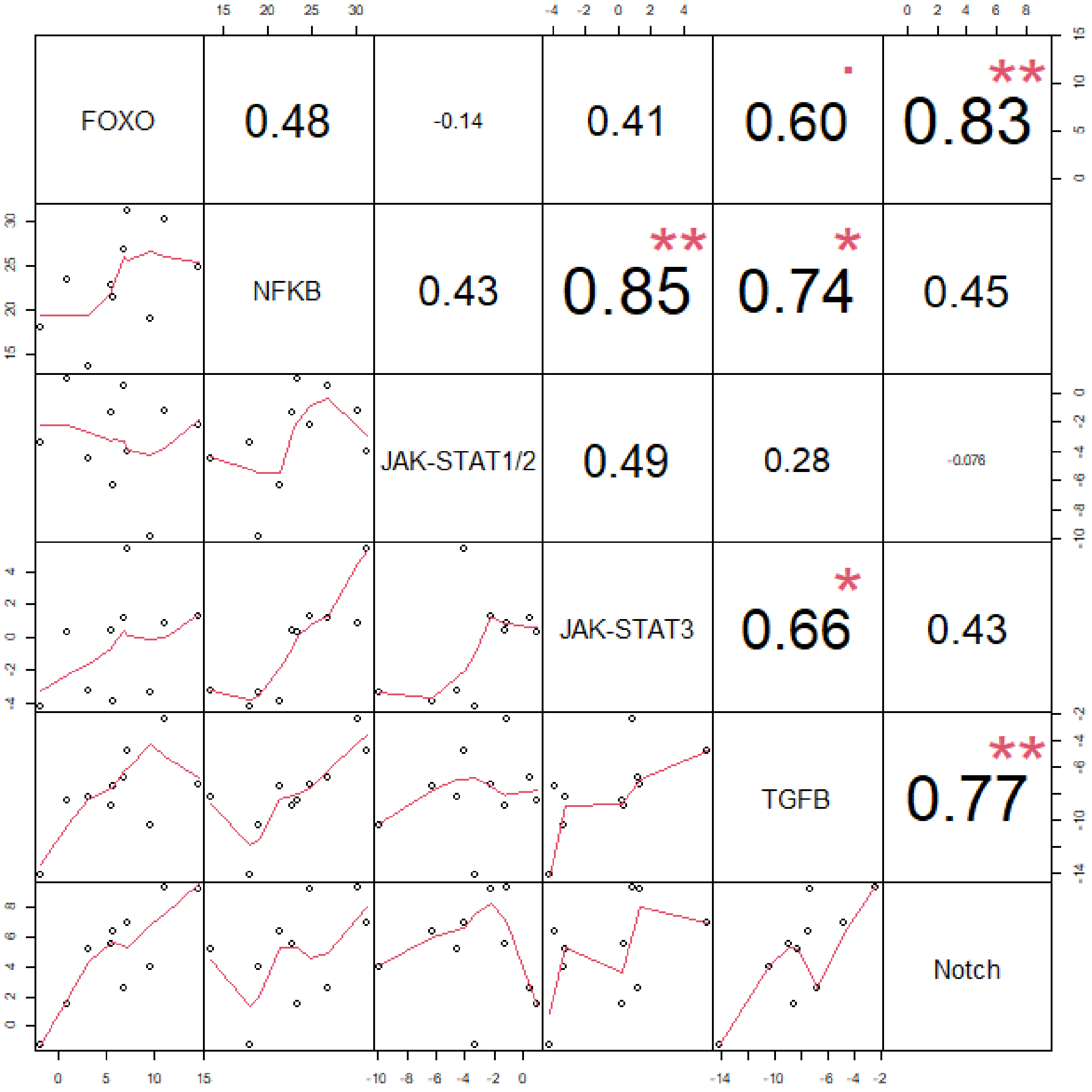


*Supplementary Figure S2. NFκB pathway activity is highly correlated with JAK-STAT3 and TGFβ pathway activity. Correlation matrix between signaling pathway activities among BrCa TIL samples from dataset GSE36765 [17]. Bottom graphs show bivariate scatterplots with a fitted line and on top are the corresponding Pearson correlation coefficients, including starts for statistically significant correlations.*

*Supplementary Table S1. Threshold values above which signaling pathway activity may be considered abnormally high. Thresholds are calculated based on activity in CD4+ T cells in blood from the four healthy donors in the GSE36765 dataset [17] and are defined as the mean pathway activity score ± two standard deviations.*

| | **lower threshold** | **upper threshold** | **# patients exceeding** the **upper threshold** |
|---|---|---|---|
| **FOXO** | -6.9 | 7.2 | 1 (pt 1) |
| **NFκB** | -12.8 | 9.4 | 1 (pt 1) |
| **JAK-STAT1/2** | -9.2 | -5.5 | 1 (pt 9) |
| **JAK-STAT3** | -12.6 | -4.4 | 2 (pt 1, 5) |
| **TGFβ** | -23.1 | -11.9 | 2 (pt 1, 5) |
| **Notch** | -9.8 | -8.5 | 9 (pt 1-2, 4-10) |

A

| | **Pearson correlation coefficients** |
|---|---|
| **FOXO** | R = 0.23, p = 0.53 |
| **NFκB** | R = 0.0094, p = 0.98 |
| **JAK-STAT1/2** | R = 0.10, p = 0.77 |
| **JAK-STAT3** | R = 0.10, p = 0.78 |
| **TGFβ** | R = 0.28, p = 0.44 |
| **Notch** | R = 0.055, p = 0.88 |

B

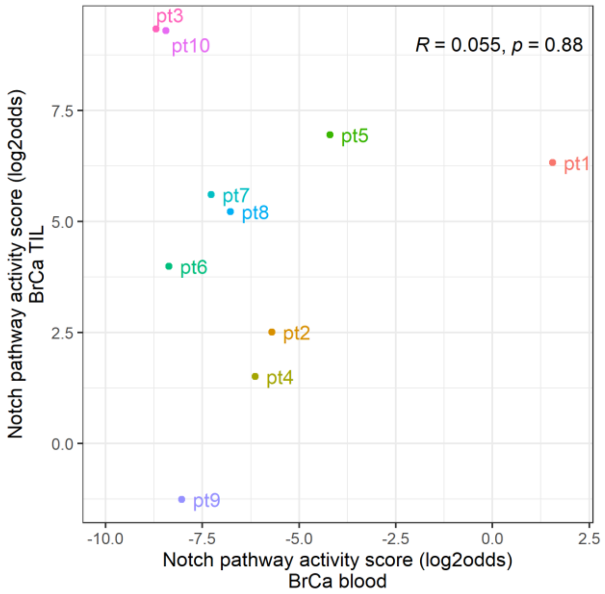


*Supplementary Figure S3. Correlation between signaling pathway activities in CD4 T+ cells from blood and from TIL from BrCa patients (dataset GSE36765, [17]). A. Pearson correlation coefficients; B. Correlation plot for the Notch pathway.*
